# Flynotyper 2.0: An updated tool for rapid quantitative assessment of *Drosophila* eye phenotypes

**DOI:** 10.1101/2024.04.07.588481

**Authors:** Johnathan Ray, Deepro Banerjee, Qingyu Wang, Santhosh Girirajan

## Abstract

About two-thirds of the genes in the *Drosophila melanogaster* genome are also involved in its eye development, making the *Drosophila* eye an ideal system for genetic studies. We previously developed Flynotyper, a software that uses image processing operations to identify and quantify the degree of roughness by measuring disorderliness of ommatidial arrangement in the fly eye. This software has enabled researchers to quantify morphological defects of thousands of eye images caused by genetic perturbations. Here, we present Flynotyper 2.0, a software that has an updated computer vision library, improved performance, and a streamlined pipeline for high-throughput analysis of multiple eye images. We also tested several batches of *Drosophila* eye images to ensure robustness and reproducibility of the updated Flynotyper 2.0 software.

**Availability and implementation:** The source code for Flynotyper 2.0 can be downloaded and installed from https://github.com/girirajanlab/flynotyper-desktop-application.

## Introduction

*Drosophila melanogaster* remains a robust model organism for genetic studies, offering valuable insights into fundamental biological processes (Reiter *et al*. 2001; Chow and Reiter 2017; Sun *et al*. 2024). As two-thirds of the genes in the *Drosophila* genome are also involved in its eye development, it stands out as an exquisite system for genetic screening and molecular studies due to its non-essential nature, ease of phenotyping, and involvement in a majority of essential genes for its development (Thaker and Kankel 1992; Thomas and Wassarman 1999; Kumar 2018). We developed Flynotyper to automatically detect and accurately quantify morphological changes observed in the rough eyes of *Drosophila melanogaster*, thus facilitating phenotypic assessment of the individual units or ommatidia of the fly eye following genetic perturbation. Given a bright-field or scanning electron microscopy derived image of the fly eye, Flynotyper uses computer vision to analyze and calculate “phenotypic scores” related to the eye’s ommatidial disorderliness. We previously demonstrated its efficacy by analyzing morphological defects resulting from the knockdown of *Drosophila* orthologs of twelve genes linked to neurodevelopmental disorders and validated through examination of eye images from six independent studies, including screens for modifiers of neurotoxicity and interactors of various genes. Its quantitative analysis accurately classified genetic modifiers of *sine oculis* obtained from genome-wide screens, demonstrating its effectiveness in assessing diverse genetic influences on eye phenotypes (Iyer *et al*. 2016). The current release of the software uses an outdated version of its computer vision tool, introducing bugs that make it difficult to use. Additionally, it can only analyze a single image per execution, provides solely a command line interface (CLI), and ran inconsistently on Mac operating systems.

Our manuscript introduces an updated version of Flynotyper that uses a modern computer vision tool supported by computational researchers and developers, providing numerous bug fixes. Additionally, it provides quality of life improvements by allowing multiple images to be analyzed at once while optimizing time spent performing such computations. Researchers can also analyze their images using a graphical-user interface (GUI). Finally, we fixed several compatibility issues that Mac users ran into. We hope that, by updating Flynotyper, it can serve as either an introduction to those who want to facilitate their fly ommatidia analysis or as an upgrade to those who are already familiar with it.

## Methods

The original version of Flynotyper utilizes OpenCV, a computer vision library. With it, the program highlights the fly eye by applying morphological transformation to the image and isolates each ommatidium with a single circle. It then goes through each ommatidium and draws six vectors from its center to the center of each bordering ommatidia. (**Figure 1**). Once all of this is done, it calculates five different values based on the distance and angle between each of the ommatidium: the total distance ommatidial disorderliness index of all stable ommatidia (ODI_D_), the total angle ommatidial disorderliness index of all stable ommatidia (ODI_A_), the total ommatidial disorderliness index of all stable ommatidia (ODI), the number of detected ommatidia (Z), and the phenotypic score (P). ODI_D_ is the difference in lengths between the five longest vectors v_1-5_ and the shortest vector v_min_, and ODI_A_ is the difference between the five largest angles formed between the vectors and the smallest angle between vectors. With these two values, ODI is calculated using their sum and the number of most ordered ommatidia N (as determined by the user). ODI and Z are then used to calculate P, which represents the severity of the eye phenotype (Iyer *et al*. 2016).

**Figure 1:**
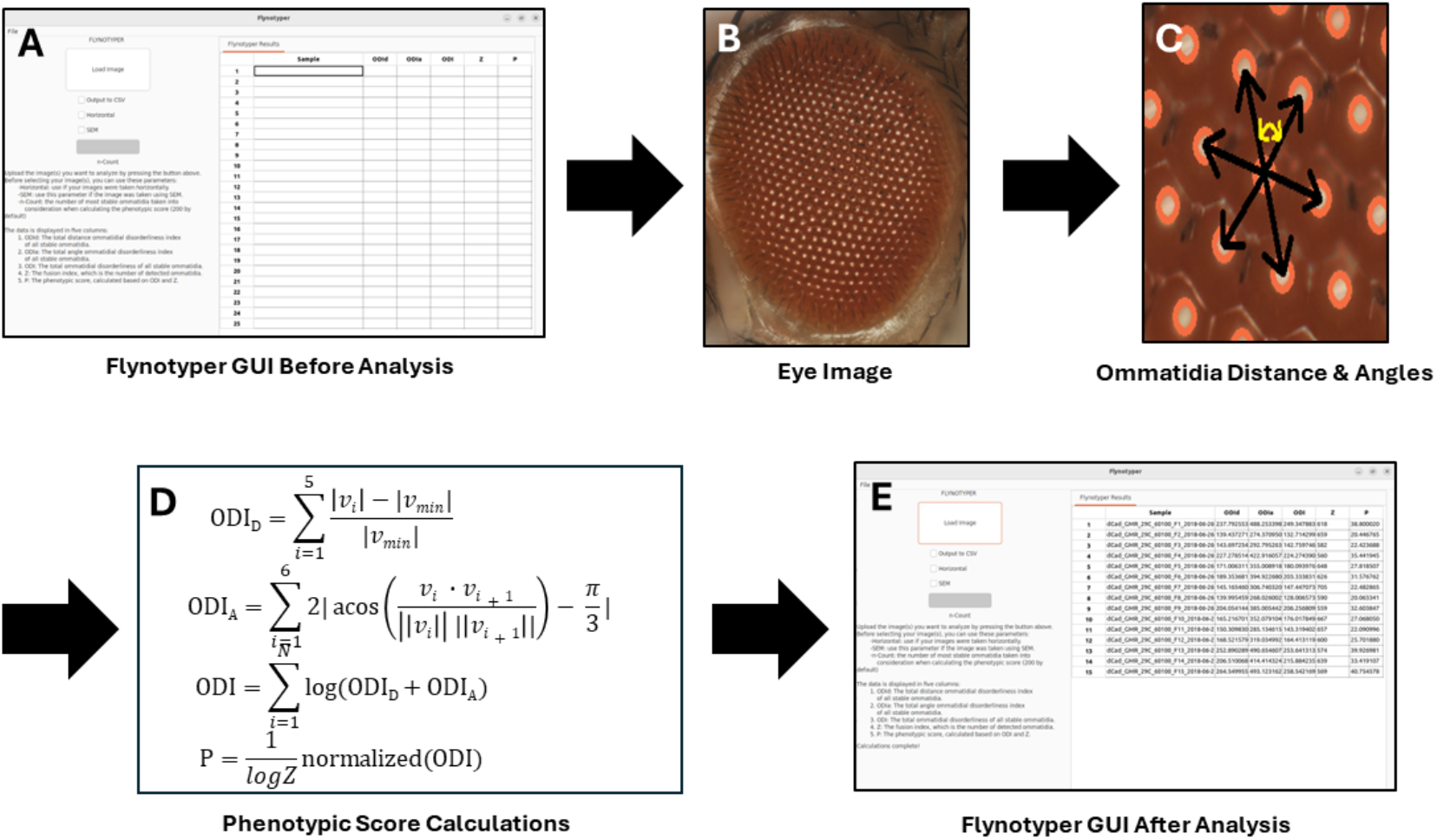
A flowchart representing the process Flynotyper 2.0 goes through to calculate its phenotypic scores and output them to the GUI. Before analysis begins, researchers are presented with a prompt to submit their images (**A**). For each image they submit (**B**), OpenCV detects and draws a circle on top of each ommatidium. It then calculates the distance from and angles between each of them (represented with the vectors found in image (**C**)). For each ommatidium, ommatidial disorderliness indices are calculated using the lengths of each of the vectors: the smallest vector, v_min_, and the five local vectors associated with it, v_i_ (i = 1…5). From there, the phenotypic score is calculated (**D**). Once this is done for all the ommatidia, the results are brought to the GUI and shown in a table (**E**).

Previously, OpenCV-2 was the most used version of the library, and we used it when creating Flynotyper. However, developers now use a different version of this library: OpenCV-4. This discrepancy in versions caused issues when installing Flynotyper. Its scripts specifically reference the OpenCV-2 library and search for binaries associated with it. If one were to install OpenCV-4, Flynotyper would not search for those binaries and assume that the researcher does not have OpenCV installed. Flynotyper 2.0 uses OpenCV-4, providing up-to-date source code for it to use. As a result, the software has fewer bugs and researchers can now install the latest version of OpenCV without having to spend time fixing issues.

In addition to analyzing images with an updated library, Flynotyper 2.0 allows researchers to run calculations on multiple images. Previously, one could only provide a single image as a command line argument for the software to analyze. This allowed results for that image to generate quickly, but at the cost of having to spend extra time rerunning the command for each individual image. Changing the source code to allow Flynotyper 2.0 to accept multiple images eliminates this. However, the software would run analysis on the images iteratively. This meant that a given image would have to wait for all the images before it to finish their calculations, resulting in more time being spent. To eliminate this added time, Flynotyper 2.0 utilizes OpenMP. This C++ library introduces parallel computing, a type of computation that allows for multiple images to be analyzed at the same time using separate CPU cores.

We also ensured that Mac users could use Flynotyper 2.0 without facing any compatibility issues. Previously, when running Flynotyper and early builds of Flynotyper 2.0, Mac systems could not find certain libraries that were required for the software to run. We realized that this was due to the way Linux and Mac had access to these libraries. For example, Linux could access OpenMP by default, whereas Mac could not. To ensure that users no longer ran into these issues, we used two different compilers based on the machine the user used: GCC for Linux and Clang for Mac.

Flynotyper 2.0 also utilizes wxWidgets, a C++ library used for creating desktop applications. With it, we created a GUI that is easy for researchers to use and understand. It provides the same functionality as its CLI counterpart while also providing some unique features of its own. Rather than passing a single image to the application, one can pass in multiple in PNG, JPG, JPEG, BMP, TIF, or TIFF formats.

Both the GUI and the CLI use a make file that researchers should run first to create the executables needed to run the software. Once this is done, they can run the executable and pass in their images. They can also use three optional flags: the horizontal flag, the SEM flag, and the count flag. By default, Flynotyper 2.0 assumes that images submitted to it are oriented vertically. The horizontal flag (-h) should be used if the images submitted are oriented horizontally. The SEM flag (-sem) is used for images that were taken with a bright-field or scanning electron microscope. The count flag (-n) updates the number of stable ommatidia taken into consideration when calculating the phenotypic score. This flag is set to 200 by default.

The source code for Flynotyper 2.0 can be installed on Mac and Linux for free here: https://github.com/girirajanlab/flynotyper-desktop-application.

## Results and Discussion

The GUI for Flynotyper 2.0 (**Figure 2**) has a similar workflow to its CLI counterpart with a few additions. Prior to uploading their images, researchers can check boxes to indicate if they would like to use the horizontal or SEM flags, and they can enter a value to update the count flag to what they would like. There is also an “Output to CSV” flag. This exports Flynotyper 2.0’s output to a CSV file format should a researcher want a record of their results. Once their images are uploaded (either as a file or a batch of them), the software conducts its calculations (**Figure 1**). For each ommatidium in the image, Flynotyper 2.0 calculates the distance between each adjacent ommatidium and angles between each of them. These values are used to calculate the phenotypic scores (**Figure 1D**). With the help of the OpenMP library, this takes minimal time. Once it is finished, the results are shown in a table. Each row contains six values: the name of the file and the five phenotypic values (ODI_D_, ODI_A_, ODI, Z, and P).

**Figure 2:**
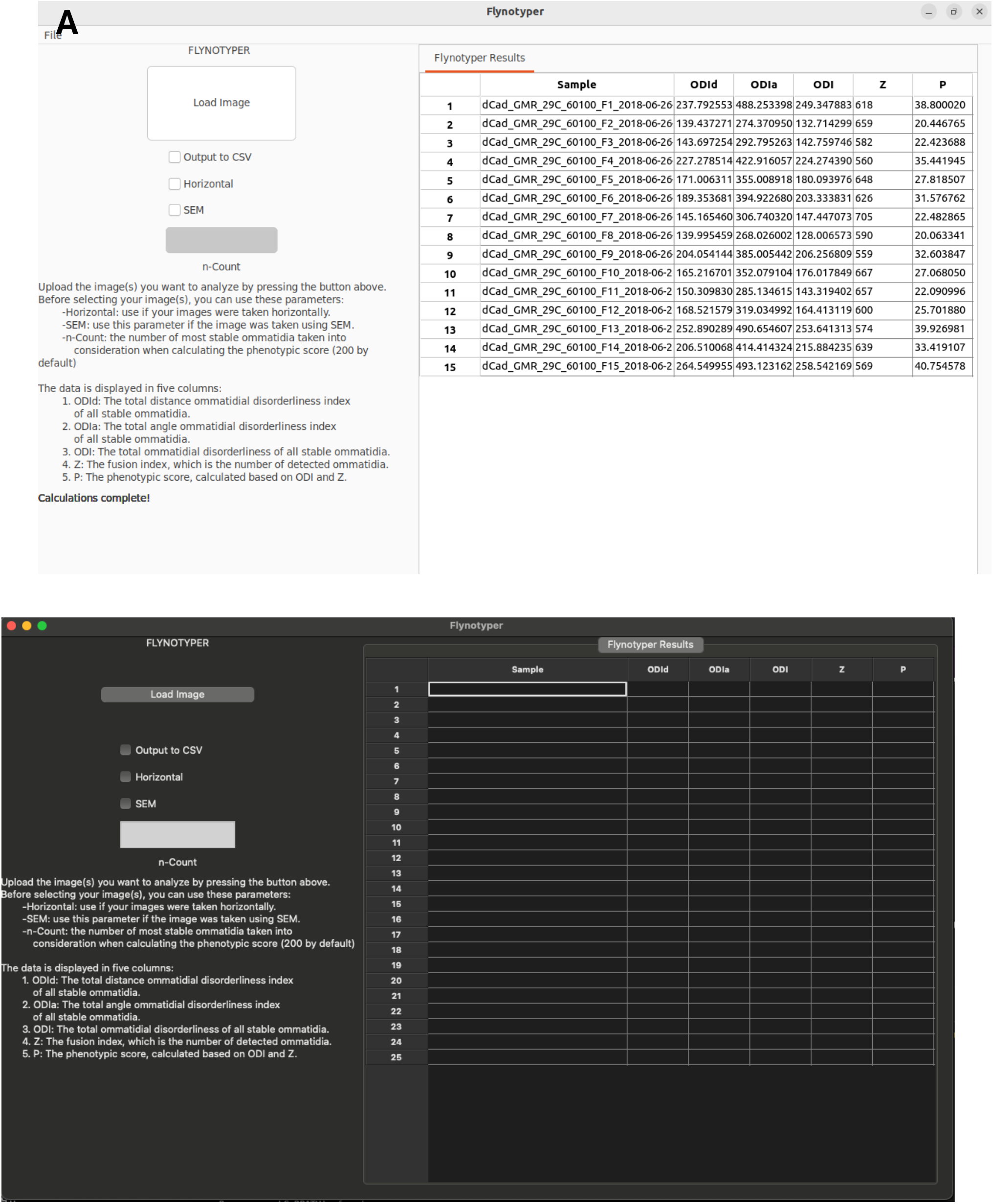
A visual of what the Flynotyper 2.0 GUI looks like on Linux (**A**) and Mac (**B**). Researchers can customize the flags they want to use prior to submitting files and select any number of images that they need analyzed. Once analysis is done, the results are displayed in a table on the right side, showing a different set of values for each image. If the “Output to CSV” flag is selected, the results are exported to a CSV file format that can be found in the directory that Flynotyper 2.0 was compiled in.

We tested the effectiveness of Flynotyper 2.0 by running analyses on four sets of images used in a previously published study which used Flynotyper to analyze *Drosophila* eyes and identify developmental, cellular, and neuronal phenotypes in genes related to the 3q29 deletion region (Singh *et al*. 2020). We found that Flynotyper 2.0 calculated values (i.e. ommatidial disorderliness indices, phenotypic scores) whose averages matched those found in the study. Additionally, to test how quickly Flynotyper 2.0 can analyze multiple images using parallel computing, we ran five separate time trials. For each trial, we used the software on the datasets twice: once without OpenMP and once with OpenMP. We ran this test on both Linux and Mac to ensure that OpenMP ran as intended on both platforms. On Linux, the analyses that did not use OpenMP and instead went through all the images iteratively took around forty seconds on average. Meanwhile, the analyses that used OpenMP took around twenty-two seconds on average, nearly cutting the calculation time in half. On Mac, similar results were found. Without OpenMP, analyses took twenty-five seconds on average, and with OpenMP, analyses took six seconds on average (**Figure 3**).

**Figure 3:**
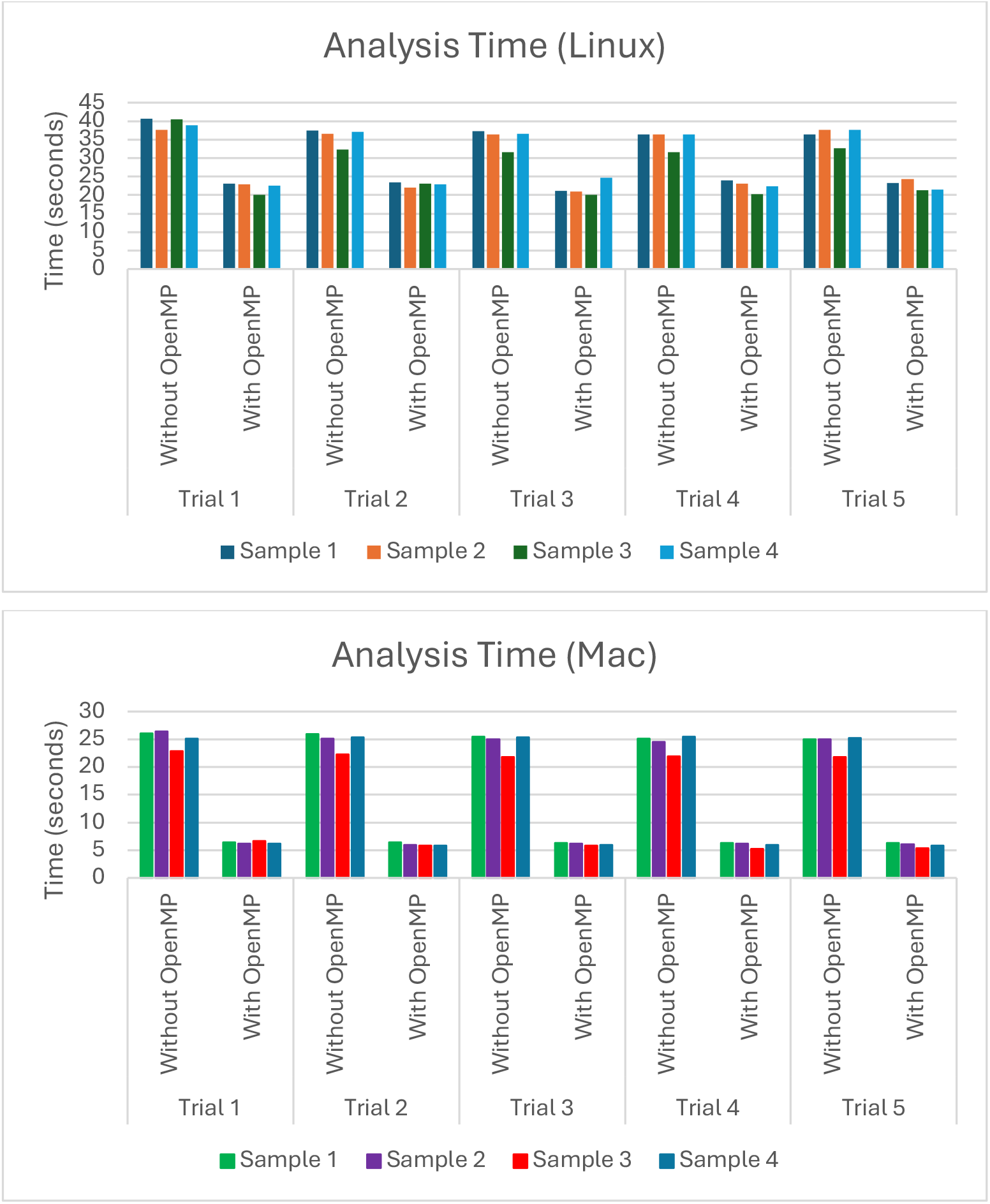
A graph showing the time it takes Flynotyper 2.0 to analyze different batches of image data on Linux and Mac. The Linux data was calculated on an Ubuntu virtual machine using an 11^th^ Gen Intel Core i5-1135G7 processor with access to two CPU cores and two gigabytes of memory. The Mac data was found using a Mac Mini with an Apple M1 processor utilizing eight cores and sixteen gigabytes of memory. On average, the time it took to analyze the images on Linux using a parallel approach was nearly half as much as doing so using an iterative approach. On Mac, it was nearly a quarter as much.

The original version of Flynotyper made numerous contributions of its own when it was first created. It can detect subtle differences in eye morphology and can quantify it. By doing so, researchers can easily see the effects such differences can have on key neurodevelopmental genes (Iyer *et al*. 2016). We hope that researchers can continue to accomplish these things in a more accessible and intuitive way using Flynotyper 2.0. The inclusion of the GUI helps users understand what each score represents, and the decreased processing time allows the user to get these scores without having to wait for the software to go through each image one at a time. With these features, one can focus on conducting effective eye phenotyping without having to put equal focus on what the software is doing. While these features do lead to an improvement in the analysis of fly eyes, Flynotyper 2.0’s effectiveness does rely on the computation power of the computer being used. Figure 3 shows that, while parallel computing does cut down analysis time dramatically, the CPU being used affects the amount of time cut down. However, we feel that this setback is small when compared to Flynotyper 2.0’s strengths.

## Conclusion

Flynotyper 2.0 is a cross-platform software that facilitates the analysis of *Drosophila* ommatidia. It provides all the benefits of its predecessor while also providing some updates of its own, including minimal software bugs, faster analysis time, the ability to input multiple images, and a GUI that visualizes the output for the researcher. We hope that this software will make the process of *Drosophila* eye phenotyping easier for those who have and have not used Flynotyper in the past.

## Data Availability Section

Fly ommatidia images are available upon request. The images used in Figure 2 are available in the “example” directory on the Flynotyper 2.0 GitHub page: https://github.com/girirajanlab/flynotyper-desktop-application/tree/main/example.

## Acknowledgements

This work was supported by the NIH grants R01-GM121907 to SG.

